# Adventitious roots facilitate surface water uptake but only partially sustain transpiration under waterlogging in tomato (*Solanum lycopersicum*)

**DOI:** 10.64898/2026.04.09.717333

**Authors:** Hermann Prodjinoto, Dor Batat, Ido Nir, Dana Menkes, Moshe Shenker, Menachem Moshelion

## Abstract

waterlogging constrains terrestrial plants by limiting oxygen diffusion in the rhizosphere and altering root-zone physical and chemical properties. However, the extent to which whole-plant responses to waterlogging can be reproduced by oxygen deficiency alone remains unresolved. In tomato (*Solanum lycopersicum*), waterlogging is commonly associated with adventitious-root formation, yet the functional contribution of these roots to whole-plant water relations has rarely been quantified.

Here, we experimentally separated root-zone hypoxia from waterlogging and quantified the contribution of surface-associated adventitious roots to whole-plant transpiration. Using high-resolution gravimetric lysimeters, we monitored transpiration dynamics under two conditions: (i) N_2_-driven displacement of root-zone O_2_ under near-field-capacity conditions and (ii) root-zone waterlogging. These measurements were complemented by analyses of soil redox potential and pH, mineral composition, stem anatomy, and genotypic variation among M82, IL11-4, and IL8-1.

N_2_-driven oxygen depletion rapidly reduced rhizosphere O_2_ concentration and induced a moderate decline in redox potential, accompanied by changes in rhizosphere chemistry and mineral relations. Whole-plant transpiration, however, declined only progressively over several days. Under waterlogging, transpiration declined rapidly in all genotypes, with strong genotype dependence. A transient partial recovery coincided with the appearance of adventitious roots at the soil surface and was followed by renewed decline after drainage.

Quantitative analysis indicated that adventitious roots contributed only a limited fraction of daily water uptake, approximately 15% to 20%, which was insufficient to restore pre-waterlogging transpiration or growth. Together, these results show that waterlogging responses were not reproduced by rapid oxygen deprivation alone and that adventitious roots provide limited hydraulic compensation.

## Introduction

Plant transpiration drives the transport of water and minerals to photosynthetic tissues. Despite the requirement for soil water to sustain this process, excess water in the rhizosphere, through root-zone flooding or waterlogging, can disrupt water and mineral uptake and thereby impair or even kill terrestrial plants. This threat is being amplified by climate change, which is increasing the frequency and intensity of heavy rainfall and flooding events. From 2000 to 2019, the global cropland area exposed to flooding increased by nearly 84,000 km^2^, equivalent to 7.75% (Zhang et al., 2023). Poorly drained regions across more than 188 countries are already experiencing more frequent and/or more severe flooding, with substantial consequences for agriculture (Rentschler et al., 2022). Currently, 24% of global cropland lies in flood-prone regions (Zhang et al., 2023), placing nearly one-quarter of global food production at recurring risk. Between 2003 and 2013, floods caused approximately US$ 7.8 billion in damage and losses to the crop subsector, making excess water the most damaging natural hazard for crops, exceeding the impacts of storms and drought (FAO, 2015).

Waterlogging rapidly displaces air from soil pores and severely restricts gas exchange between the soil and the atmosphere. As pore spaces become filled with water, gas diffusion declines by several orders of magnitude, sharply reducing oxygen availability to roots (Yalin et al., 2021). Oxygen deficiency, widely considered a primary cause of flooding injury in terrestrial plants, shifts root metabolism from aerobic respiration to anaerobic pathways, thereby constraining root respiration, reducing metabolic activity, and creating energy deficits that lead to progressive root damage (Colmer and Voesenek, 2009; Horchani et al., 2009; Pan et al., 2021). In tomato, flooded soils typically induce a sequence of symptoms that includes leaf wilting, strong epinasty of intermediate leaves, reduced shoot elongation, callus formation, adventitious-root emergence along the stem near the water surface, and chlorosis of older leaves (Jackson, 1956). Flooding also impairs transpiration and growth, reducing stem elongation and altering shoot physiology (Kuo and Chen, 1980). Similar reductions in carbon exchange and transpiration have also been reported in other species under water excess, including pigeonpea (Bansal and Srivastava, 2015).

In addition to limiting oxygen supply, flooding alters rhizosphere chemistry. Low-oxygen conditions promote the accumulation of gases such as CO_2_ and methane and stimulate anaerobic microbial processes, beginning with denitrification and, under more reducing conditions, progressing to the reduction of manganese, iron, and sulfate. These processes alter soil redox potential (Eh) and can strongly affect nutrient availability to plants (Colmer and Voesenek, 2009; Pezeshki and DeLaune, 2012; Singh et al., 2018). Consequently, nutrient uptake and transport may be impaired, potentially leading to deficiency or toxicity despite ongoing plant acclimation. For example, 2-7 days of flooding in barley reduced foliar N, P, and K concentrations by 50%-60%, illustrating how water excess can impair nutrient acquisition through both altered soil chemistry and oxygen-limited root function (Leyshon and Sheard, 1974). Nitrogen, phosphorus, and potassium are among the nutrients most strongly affected by rhizosphere water excess in terrestrial plants (Aslam et al., 2023).

Accordingly, we monitored rhizosphere pH and redox potential, characterized elemental dynamics in the soil solution and drainage, and quantified mineral composition in leaves and roots to determine whether oxygen depletion induces rhizosphere redox changes that influence metal dynamics and whole-plant nutrient status.

Under waterlogging (often referred to as flooding), many terrestrial plants develop adventitious roots, typically emerging from the stem or lower shoot near the soil or water surface (Jackson, 1956; Sauter, 2013). These roots are commonly associated with flooded environments and have been widely linked to oxygen deficiency and ethylene signaling in the submerged root zone (McNamara and Mitchell, 1990; Vidoz et al., 2010; Kęska et al., 2021). However, adventitious-root formation occurs specifically under waterlogged conditions, where roots are exposed to free or near-surface water, raising the possibility that its induction is not driven solely by hypoxia but also by the distinct hydraulic and physical configuration imposed by water excess. Unlike the primary root system embedded in soil, adventitious roots often develop at the soil-air or soil-water interface, where oxygen availability and water accessibility differ fundamentally from those under hypoxia alone. Although adventitious roots are often interpreted as an adaptive response that replaces or supplements the primary root system under flooding (Colmer and Voesenek, 2009; Sauter and Steffens, 2014), their actual contribution to whole-plant water uptake and transpiration has rarely been quantified.

Despite extensive documentation of adventitious-root formation under flooding, two major gaps remain unresolved. First, most studies implicitly attribute flooding responses to oxygen deficiency alone, without experimentally separating hypoxia from the physical conditions created by water excess, in which roots are exposed to free surface water rather than soil. It therefore remains unclear whether key physiological responses to flooding, including changes in transpiration and nutrient uptake, can be explained solely by oxygen limitation or instead reflect additional water-excess-specific constraints on whole-plant water relations. Second, although adventitious roots are widely interpreted as a functional replacement for the primary root system under flooded conditions, their hydraulic contribution at the whole-plant level has rarely been quantified. In particular, direct links among adventitious-root development, surface-water uptake, and whole-plant transpiration remain largely inferential, limiting our ability to determine whether these roots actively sustain plant water balance or mainly indicate the severity of flooding-induced stress.

To address these questions, we first examined the response of tomato cv. M82 to oxygen deficiency imposed by N_2_ displacement of soil oxygen, focusing on whole-plant transpiration dynamics and mineral uptake. We then compared these responses with those of two tomato introgression lines, IL11-4 and IL8-1, previously shown to differ in whole-plant transpiration relative to the recurrent parent M82. Because these genotypes differ in transpiration capacity, they provide a useful framework for testing whether baseline hydraulic and stomatal behavior modulates the extent to which adventitious roots sustain water uptake and transpiration under waterlogging. Together, these experiments were designed to disentangle the effects of root-zone oxygen limitation from the water-excess configuration imposed by waterlogging, and to test whether adventitious-root development provides a functional pathway for surface-water uptake that contributes to sustaining whole-plant transpiration. Specifically, we asked whether waterlogging responses can be explained solely by reduced rhizosphere oxygen availability, approximated here by N_2_-induced O_2_ displacement under otherwise normal irrigation at field capacity, or whether water excess imposes additional constraints on whole-plant water relations that cannot be inferred from oxygen deprivation alone. We further asked how hypoxia and waterlogging affect whole-plant transpiration dynamics, mineral relations, and development, and how rapidly these responses can be detected using high-resolution time-series measurements. Finally, we asked to what extent adventitious roots contribute to surface-water uptake and whole-plant transpiration under waterlogging, and whether genotypes with contrasting transpiration capacity differ in this response. Based on prior literature, we hypothesized that (H1) root-zone oxygen deficiency constrains root respiration, thereby reducing whole-plant transpiration and altering mineral uptake, and that (H2) under waterlogging, adventitious roots make a measurable contribution to surface-water uptake that partially offsets transpiration losses, whereas under N_2_-induced hypoxia their hydraulic contribution is minimal.

## Results

Our initial experiments examined whole-plant transpiration responses to root-zone oxygen deficiency imposed independently of soil water excess.

### Oxygen deficiency induced by O_2_ displacement (non-flood hypoxia)

#### Effect of oxygen level on transpiration dynamics, soil pH, and redox potential

Following the onset of N_2_ injection (Figure 3, red dashed line), soil O_2_ concentration in the oxygen-deficient treatment declined rapidly, reaching approximately 1.7% within 1.5 h and stabilizing at 0.09%-0.7%, whereas ventilated controls remained close to pre-treatment levels (Figure 3B; Supplementary Figure S3B). Despite this rapid O_2_ depletion, midday whole-plant transpiration did not differ significantly between treatments during the first approximately 20 h. A significant reduction in transpiration under oxygen deficiency developed only after sustained hypoxia, resulting in a mean decrease of approximately 21%-22% during days 5-10 of treatment in Exp. 2 and days 2-6 in Exp. 1 (Figure 3A; Supplementary Figure S3A). This temporal lag indicates that the transpiration response was not an immediate consequence of O_2_ depletion, but rather reflected downstream physiological adjustment to prolonged hypoxic conditions.

**Figure 1.**
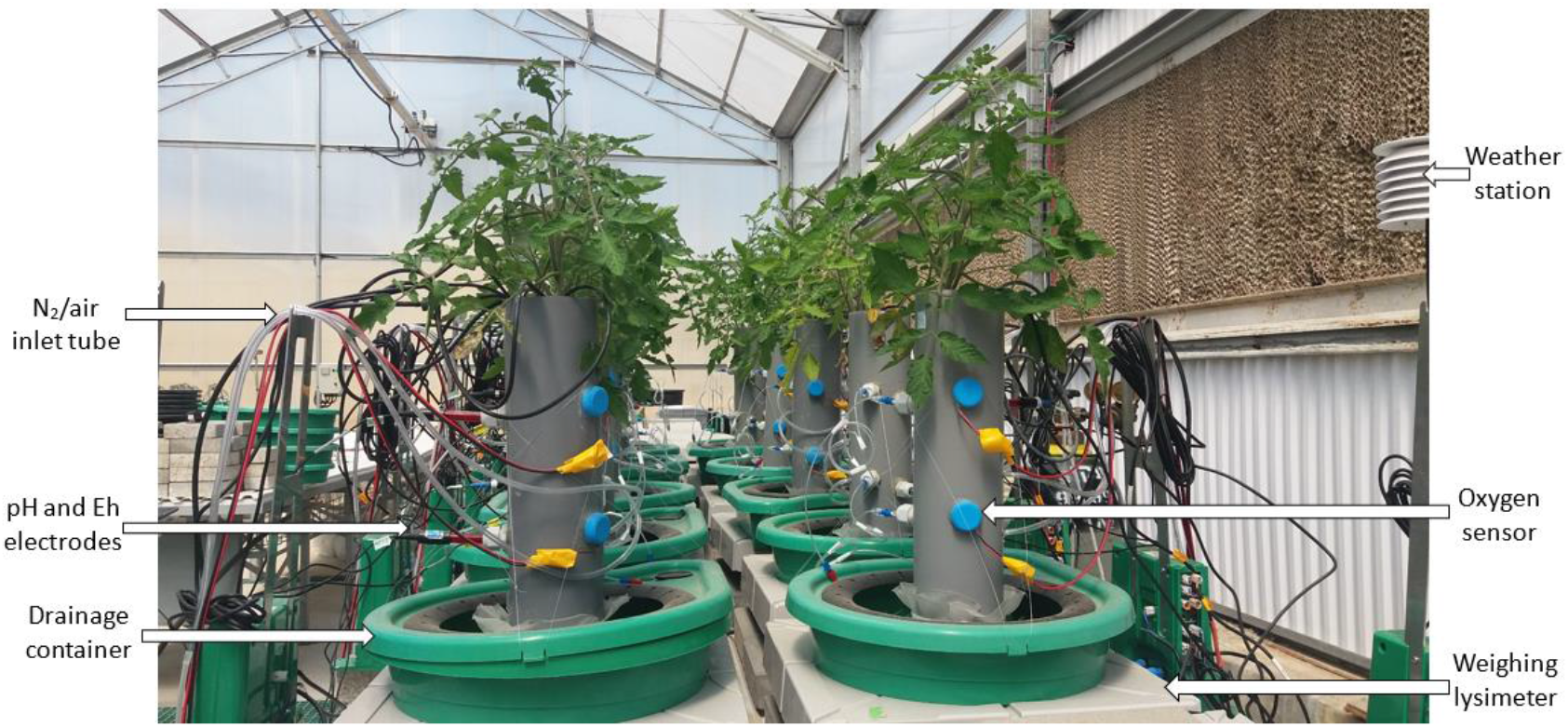
Experimental platform used to impose non-flood hypoxia while continuously monitoring whole-plant and rhizosphere responses. Tomato plants were grown in PVC soil columns mounted on weighing lysimeters (PlantArray 3.0) positioned above drainage containers. Hypoxic or aerated conditions were imposed by continuous delivery of N_2_ or air through an inlet tube inserted into the soil column. Rhizosphere pH and redox potential (Eh) were monitored using electrodes inserted through dedicated side ports, and soil oxygen was measured using an oxygen sensor installed in the column wall. Environmental conditions were recorded by the greenhouse weather station. The system enabled continuous gravimetric, environmental, and rhizosphere monitoring throughout the experiment.

**Figure 2.**
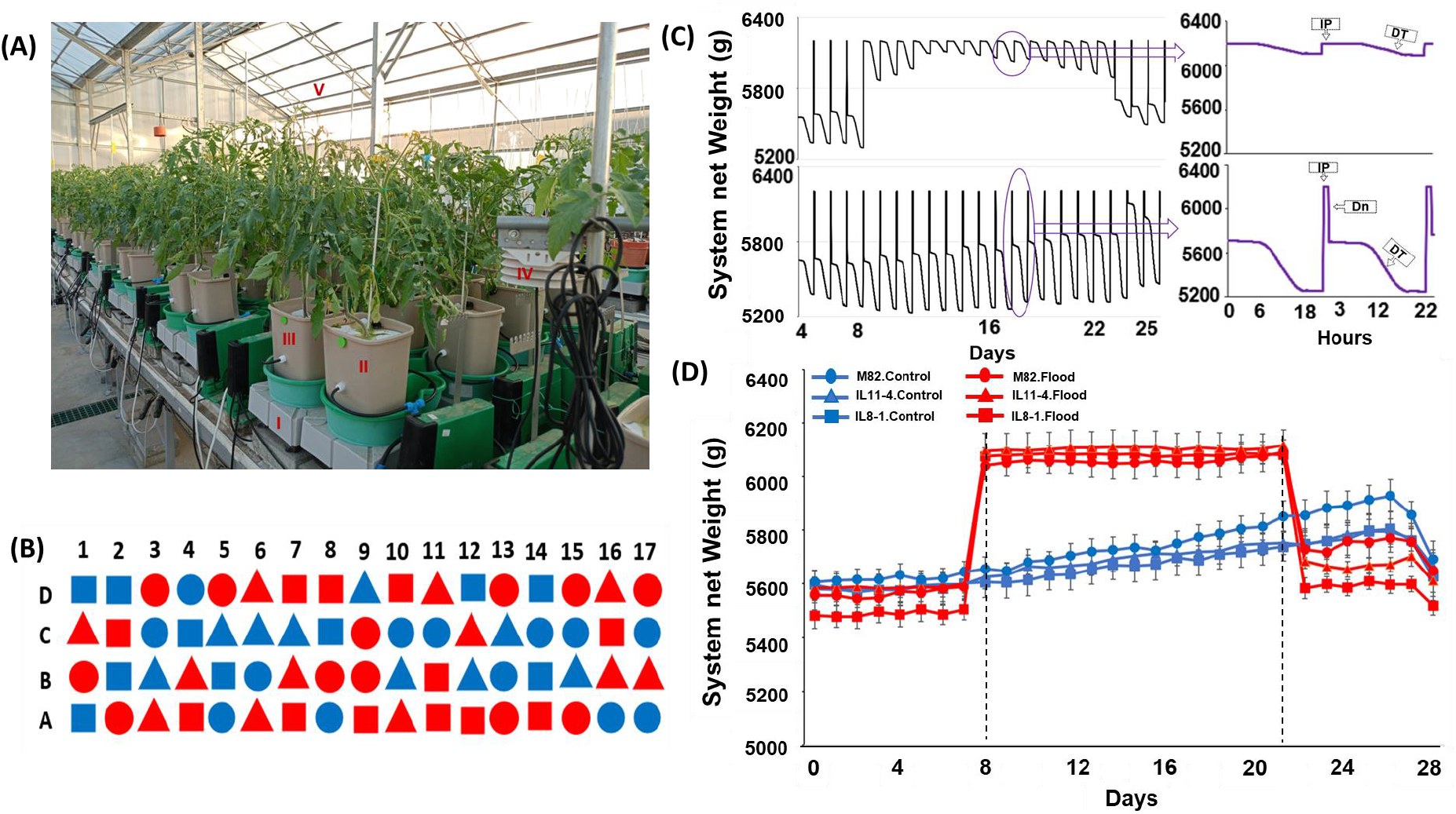
Waterlogging experimental setup and design. (A) Photograph of the waterlogging experiment conducted on the PlantArray gravimetric phenotyping platform, showing: (I) lysimeter platform, (II) waterlogged pot, (III) control pot, (IV) greenhouse weather station, and (V) polycarbonate greenhouse. (B) Randomized spatial arrangement of the 68 pots on the platform. Shapes indicate genotypes (M82, circle; IL11-4, triangle; IL8-1, square). Blue symbols denote control plants, and red symbols denote waterlogged plants. (C) Representative system-weight traces illustrating waterlogging (top) versus standard irrigation-drainage cycles (bottom). Changes in system net weight in the waterlogged pot primarily reflect whole-plant transpiration, as drainage is prevented and soil evaporation is minimized by the EVA surface cover. IP, irrigation peak; Dn, drainage; DT, daily transpiration. (D) Daily pre-dawn system weight (plant, soil, and water) measured at 04:00, after drainage and before the onset of daily transpiration. Pre-dawn measurements minimize the effects of diurnal transpiration and short-term irrigation dynamics; therefore, day-to-day increases in system weight represent plant growth. Vertical dashed lines indicate the start and end of the waterlogging period. Lines represent genotype-by-treatment means, with symbols as in (B); waterlogged treatments are shown in red and control treatments in blue.

**Figure 3.**
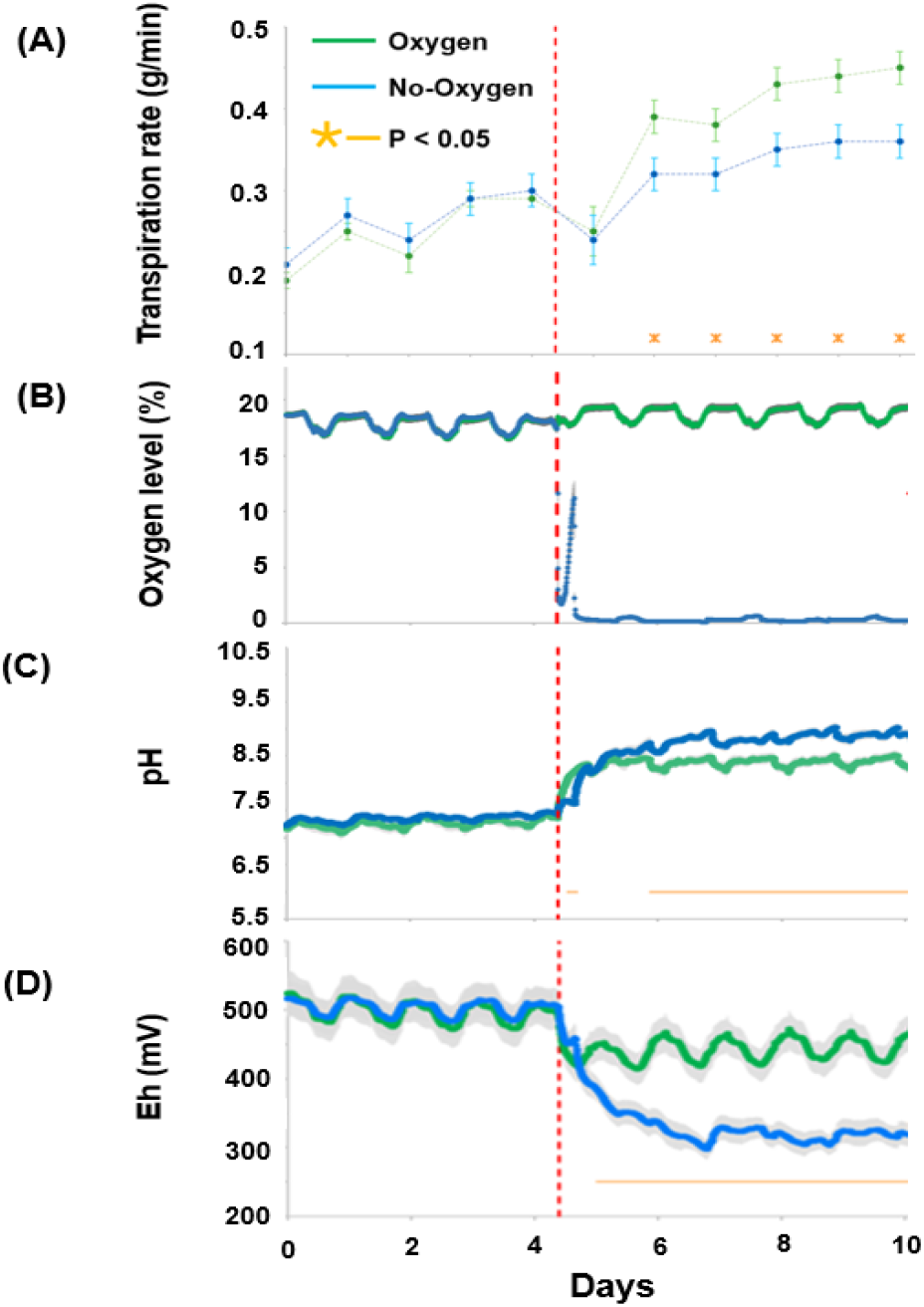
Overview of key experimental parameters in Exp. 2. Continuous time-series measurements of (A) midday whole-plant transpiration rate, (B) soil O_2_ concentration measured at 12.5 and 27.5 cm depths and presented as depth-averaged values, (C) soil pH, and (D) redox potential (Eh). The red dashed line marks the onset of N_2_ injection. Data are means ± SE. Orange symbols or horizontal bars indicate time points or intervals with significant differences between treatments (P < 0.05). Sample sizes were n = 5 for the oxygen treatment and n = 4 for the oxygen-deficiency treatment in (A), n = 9 and n = 10, respectively, in (B), n = 8 per treatment in (C), and n = 10 per treatment in (D).

O_2_ depletion was accompanied by marked changes in soil chemical conditions. Soil pH increased following O_2_ displacement by N_2_ gas flow. The increase relative to the naturally aerated treatment in Exp. 1 (mean increase of approximately 0.9 pH units; Supplementary Figure S3C) was greater than that relative to the forced-air treatment in Exp. 2 (mean difference of approximately 0.4 pH units; Figure 3C). In addition, in Exp. 2, pH increased in both treatments, whereas in Exp. 1 the difference between treatments was established within a few hours; in Exp. 2, it developed more gradually, becoming apparent only after approximately 1.5 d. Redox potential (Eh) shifted rapidly toward more reducing conditions under oxygen deficiency. Whereas ventilated controls showed only a modest decline of approximately 50 mV, Eh in oxygen-deficient columns decreased sharply within the first 15 h and continued to decline thereafter, with mean values of 445 mV in controls and 327 mV under oxygen deficiency across days 5-10 (Figure 3D; Supplementary Figure S3D). Together, these changes confirm the establishment of a more reduced soil environment under sustained O_2_ deprivation.

#### Elemental concentrations under hypoxia

Elemental responses to hypoxia were most pronounced in the soil solution and drainage, whereas changes in plant tissues were limited and element specific. Hypoxia was imposed by N_2_ injection and compared with aerated controls. In the soil solution and drainage, clear treatment effects were detected, most notably a consistent increase in potassium (K) concentration in drainage under oxygen deficiency (Figure 4A,C). Among microelements, treatment-dependent differences were also observed in both soil solution and drainage, although these responses were more element specific and less consistent than those of the macroelements (Figure 4B,D). In the youngest mature leaf, concentrations of S, Na, K, Ca, and P differed between treatments and were generally higher in ventilated controls, indicating better maintenance of leaf mineral status under aerobic conditions, with the exception of Na. Magnesium concentration did not differ between treatments (Figure 4E). In roots, only K showed a significant treatment effect and was higher under oxygen deficiency (Figure 4G), suggesting altered K accumulation or retention in root tissues under hypoxic conditions. A seasonal replicate (Exp. 1) supported these trends. Before treatment initiation, soil-solution elemental concentrations were similar between treatments (Supplementary Figure S4A,C).

**Figure 4.**
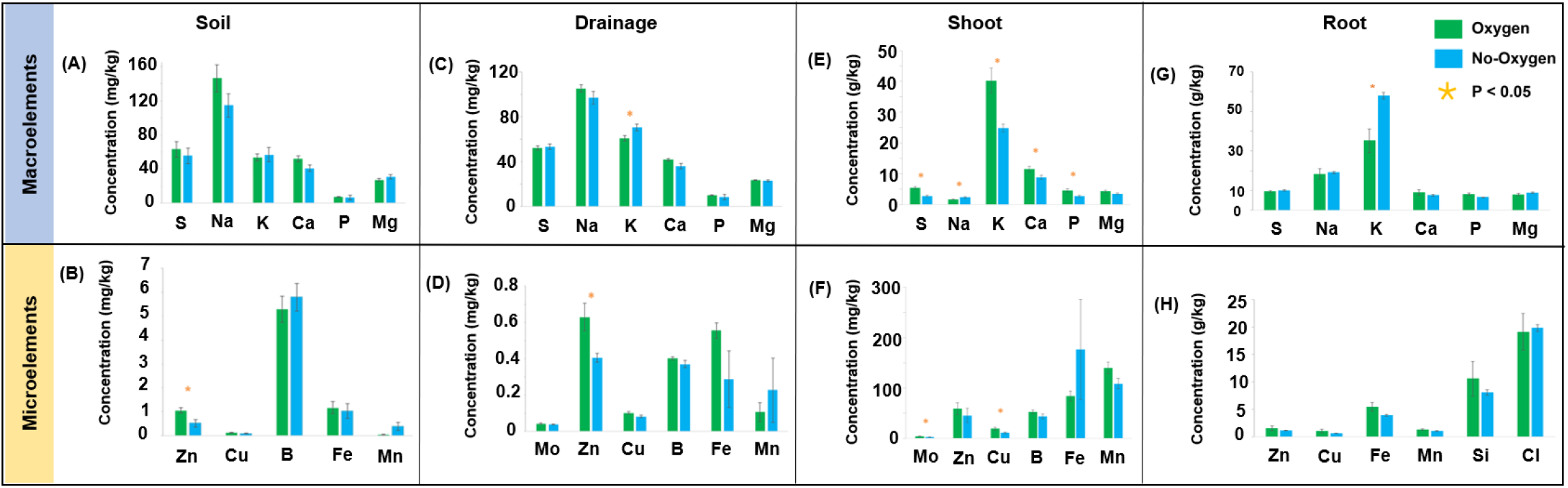
Elemental concentrations in soil solution, drainage, and plant tissues under oxygen deficiency in Exp. 2. (A-D) Elemental concentrations in soil solution and drainage water. (E,F) Elemental concentrations in the youngest mature leaf. (G,H) Elemental composition of roots determined by SEM-EDS. Data are means ± SE. Orange symbols indicate statistically significant differences between treatments for the same element (P < 0.05).

Following hypoxia, K and Mg increased in oxygen-deficient pots, whereas Mn decreased relative to controls, while most other elements showed minor or no changes (Supplementary Figure S4B,D).

Notably, although most physiological and chemical responses to hypoxia were consistent with previous reports, neither hypoxia experiment induced adventitious-root formation. This outcome contrasted with the prevailing literature and motivated a subsequent experiment in which oxygen deficiency was imposed by waterlogging rather than by N_2_-driven displacement of soil O_2_.

### Waterlogging experiment

#### Adventitious root-associated contribution to transpiration

Plants were maintained under standard irrigation-drainage cycles for 8 d, after which waterlogging was imposed by flooding the root zone on day 9. In all genotypes, midday whole-plant transpiration declined rapidly following waterlogging, reaching minimum values on days 13-14, with mean reductions of approximately 16% in M82, 44% in IL11-4, and 55% in IL8-1 relative to their respective controls (Figure 5A-C). After several days of continuous waterlogging, a transient partial recovery in transpiration was observed, reflected by a gradual rise in transpiration rates (dotted green lines), coinciding with the appearance of adventitious roots at the soil-air interface (Figure 5D-G). Adventitious-root formation was strongly genotype dependent: control plants showed few or no adventitious roots, whereas waterlogged plants exhibited significantly higher adventitious-root scores, highest in IL8-1, intermediate in IL11-4, and lowest in M82 (Figure 5H).

**Figure 5.**
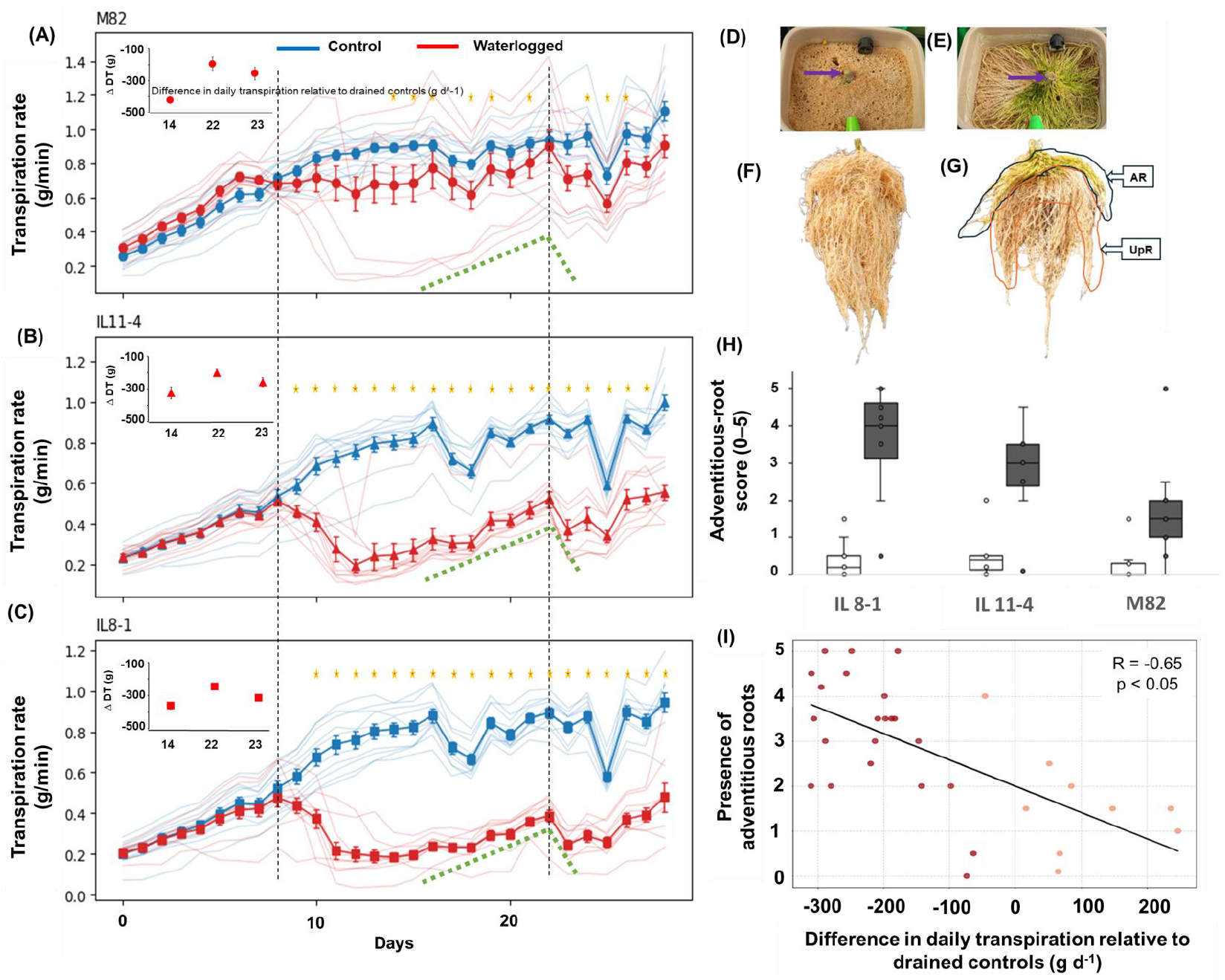
Whole-plant transpiration dynamics and adventitious-root formation under waterlogging. (A-C) Midday whole-plant transpiration rates during waterlogging in M82 (A), IL11-4 (B), and IL8-1 (C). Bold lines represent means ± SE, and thin lines represent individual plants. Blue lines indicate drained controls, and red lines indicate waterlogged plants. Vertical dashed lines indicate the onset and termination of waterlogging. Orange asterisks denote significant differences between treatments at the corresponding time points (Student’s t-test, P < 0.05). Dashed green lines indicate the period during which adventitious roots were visible at the soil-air interface and transpiration partially recovered. Insets show the difference in daily transpiration rate relative to the control (ΔDT; g day^-^1), obtained by temporal integration of transpiration data over the indicated time windows. (D) Representative surface view of a control pot showing no visible adventitious roots. (E) Representative surface view of a waterlogged pot showing adventitious roots emerging at the soil surface. (F) Corresponding whole-root system of the same plant shown in (D), imaged at harvest. (G) Representative whole-root system of a waterlogged plant at harvest, with an annotated schematic highlighting adventitious roots (AR) and upper roots (UpR). Waterlogging was associated with increased adventitious rooting and enhanced upper-root development, together with reduced development of deeper roots. (H) Adventitious-root scores at harvest (0-5 scale). Open boxes represent controls, and filled boxes represent waterlogged plants. (I) Relationship between the difference in daily transpiration rate relative to drained controls and adventitious-root score under waterlogging. Each point represents an individual waterlogged plant; dark red points: IL11-4 and IL8-1and light orange points: M82; the line indicates linear regression (Pearson r = -0.65, P < 0.05). Sample sizes: M82, n = 11 per treatment; IL11-4, n = 12 controls and n = 10 waterlogged plants; IL8-1, n = 11 controls and n = 10 waterlogged plants.

Across individual waterlogged plants, adventitious-root score was negatively correlated with the reduction in transpiration following flooding (Pearson r = -0.65; Figure 5I), such that plants exhibiting stronger adventitious-root development also showed larger absolute reductions in transpiration. This relationship suggests that adventitious-root formation tracked stress severity under waterlogging. At the same time, the temporal pattern, namely a partial recovery phase during prolonged waterlogging followed by a renewed decline after drainage, is consistent with a late, partial compensatory contribution of surface-associated roots to water uptake. Similar patterns were observed in daily transpiration dynamics (Supplementary Figure S5).

Importantly, the magnitude and consistency of transpiration decline varied among genotypes: all waterlogged IL8-1 plants exhibited reduced transpiration, all but one IL11-4 plant showed a reduction, whereas only three M82 plants displayed a clear decrease. Accordingly, quantitative analyses of transpiration losses were restricted to plants that exhibited a measurable decline in transpiration following waterlogging. On days 13-14, reductions in daily transpiration relative to controls amounted to 423.36 ± 3.82 g in M82, 324.22 ± 36.16 g in IL11-4, and 361.60 ± 21.57 g in IL8-1 (insets in Figure 5A-C), indicating genotype-specific sensitivity to waterlogging. On day 22, drainage was restored and plants returned to irrigation-drainage cycles, resulting in a renewed decline in transpiration as adventitious roots lost access to surface water (dotted green lines). During the period of partial compensation (days 14-22), mean daily transpiration losses relative to controls decreased to 196.25 ± 43.42 g in M82, 201.53 ± 21.91 g in IL11-4, and 246.65 ± 17.38 g in IL8-1. Following drainage of the surface water layer (approximately 96 g), mean daily transpiration losses increased again to 255.57 ± 40.03 g in M82, 259.36 ± 27.51 g in IL11-4, and 314.85 ± 13.73 g in IL8-1.

Based on the extent of adventitious-root coverage at the soil surface and the available surface-water volume, the effective contribution of these roots to water uptake was estimated at 59.32 g in M82, 57.83 g in IL11-4, and 68.2 g in IL8-1. Together, these estimates indicate that adventitious roots made a measurable contribution to sustaining transpiration under prolonged waterlogging, but that this contribution was insufficient to fully offset waterlogging-induced transpiration losses.

Cumulative transpiration was reduced by nearly half in waterlogged IL8-1 and IL11-4 plants relative to their drained controls, whereas M82 did not differ significantly from its control (Figure 6A). Shoot dry weight also declined by approximately half in waterlogged IL8-1 and IL11-4 plants, whereas M82 was not significantly affected (Figure 6B). Waterlogging reduced shoot water-use efficiency (WUE), calculated here as shoot dry weight per unit cumulative transpiration, in all three genotypes relative to their controls, with IL8-1 showing the lowest values (Figure 6C). Shoot length was significantly reduced only in IL8-1, with no significant effect detected in IL11-4 or M82 (Figure 6D). Root length decreased in IL8-1 and IL11-4 relative to their controls, whereas M82 showed no significant change (Figure 6E). The root:shoot ratio increased in IL8-1 and IL11-4 relative to their controls, but not in M82 (Figure 6F). Visual assessment at harvest supported these genotype-dependent morphological responses to waterlogging (Figure 6G). Under waterlogging, IL8-1 and IL11-4 appeared smaller and showed more leaf wilting than their drained controls, whereas M82 appeared less affected. Root systems of waterlogged IL8-1 and IL11-4 plants also showed browning and lower apparent root mass, whereas M82 roots appeared less affected.

**Figure 6.**
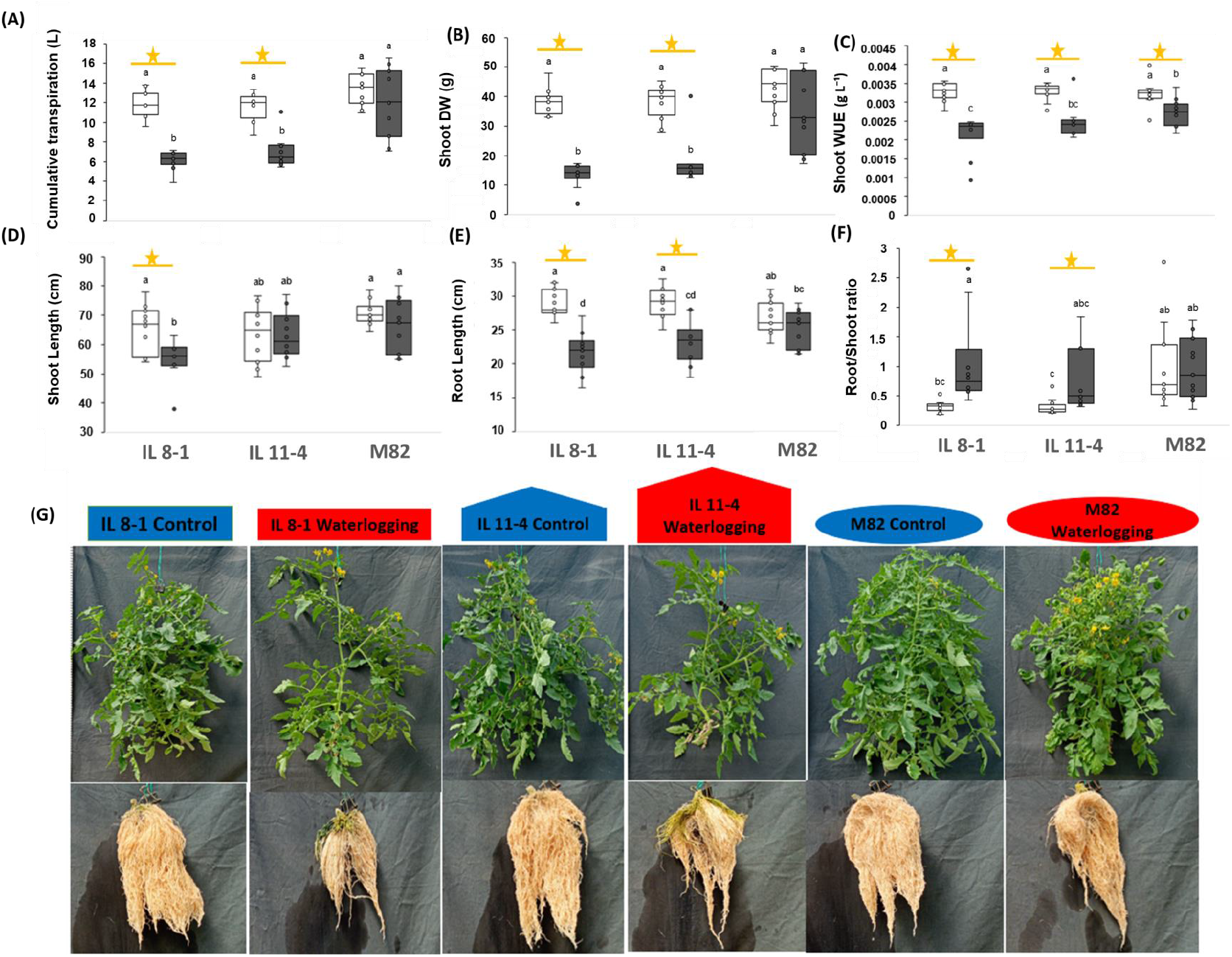
Effects of waterlogging on transpiration, biomass, water-use efficiency, and plant morphology. (A) Cumulative transpiration. (B) Shoot dry weight at harvest. (C) Shoot water-use efficiency (WUE, g L^−1^), calculated as shoot dry weight divided by cumulative transpiration. (D) Shoot length. (E) Root length. (F) Root:shoot ratio. (G) Representative shoots and corresponding root systems of IL8-1, IL11-4, and M82 plants grown under control and waterlogging conditions at harvest. Data are presented as boxplots. Open boxes represent control plants, and filled boxes represent waterlogged plants. Sample sizes: M82, n = 11 per treatment; IL11-4, n = 12 controls and n = 10 waterlogged plants; IL8-1, n = 11 controls and n = 10 waterlogged plants. Different letters indicate significant differences among genotype-by-treatment groups (one-way ANOVA followed by Tukey’s HSD, P < 0.05). Orange lines with asterisks indicate within-genotype differences between control and waterlogging treatments (Student’s t-test, P < 0.05).

To assess whether waterlogging effects extended to reproductive development, we quantified flower and fruit production across genotypes at harvest (Supplementary Figure S6). Stem cross-sections taken just above the adventitious-root zone, corresponding to the region shown in Figure 5D and E, suggested qualitative anatomical differences under waterlogging. Relative to controls, waterlogged stems showed an apparent reduction in the visually distinguishable xylem area, with this pattern appearing more pronounced in IL11-4 and IL8-1 than in M82 (Figure 7).

**Figure 7.**
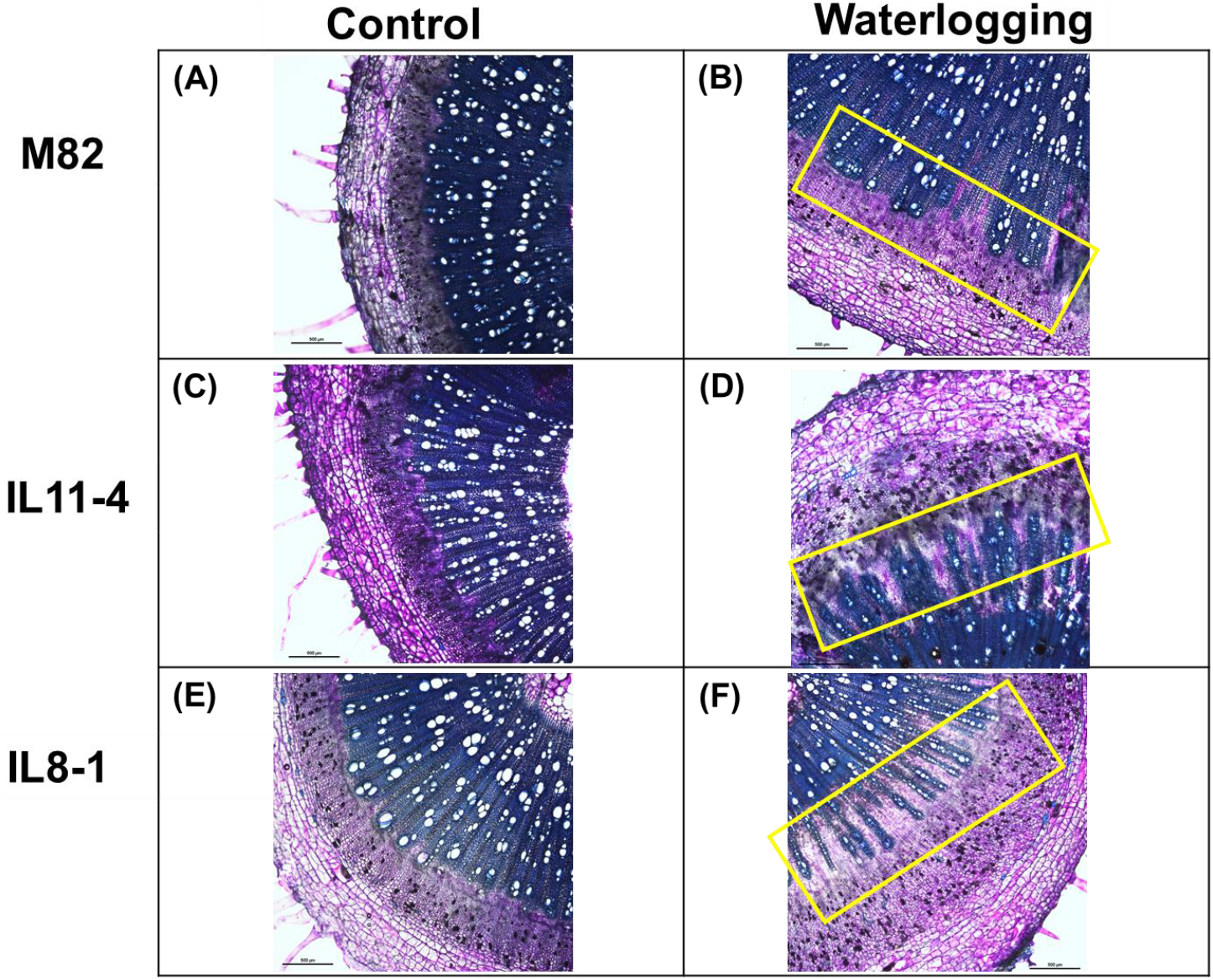
Toluidine blue-stained tomato stem cross-sections at harvest. (A,B) M82; (C,D) IL11-4; (E,F) IL8-1. Left panels show control plants, and right panels show waterlogged plants. Sections were collected just above the adventitious-root zone, stained with toluidine blue, and imaged using an Olympus BX53 microscope equipped with a DP73 camera. Yellow boxes highlight the xylem region, where visible anatomical differences were observed under waterlogging.

## Discussion

In tomato, the whole-plant response to waterlogging cannot be inferred from oxygen deprivation alone as imposed by rapid N_2_ displacement, but instead reflects the interaction between root-zone O_2_ limitation and genotype-specific differences in hydraulic performance, metabolic capacity, and developmental plasticity, consistent with conceptual frameworks of flooding responses (Voesenek and Bailey-Serres, 2015) and with physiological and signaling studies under excess water (Jia et al., 2021). Here, we disentangled root-zone O_2_ deprivation from waterlogging as a distinct hydraulic condition and tested whether adventitious roots provide a functional pathway for surface-water uptake that contributes to whole-plant water balance. We hypothesized that (H1) O_2_ displacement constrains root respiration, thereby reducing whole-plant transpiration and mineral uptake, and that (H_2_) waterlogging induces a compensatory response through adventitious roots that partially sustains transpiration, whereas under hypoxia without water excess their hydraulic contribution is negligible. Our results support a two-layer response. Hypoxia alone caused a delayed but persistent reduction in water and mineral fluxes without inducing adventitious-root formation, whereas waterlogging imposed additional physical boundary conditions that triggered adventitious-root formation and a transient partial recovery of transpiration, with strong genotype-dependent consequences for whole-plant performance.

N_2_-induced hypoxia imposed clear and reproducible physiological constraints on tomato plants. Across two independent experiments, displacement of soil O_2_ by N_2_ resulted in pronounced changes in soil chemistry, including increased pH and a marked decline in redox potential (Figure 3C,D; Supplementary Figure S3C,D), together with shifts in mineral composition in leaves and roots (Figure 4E-H). These responses are consistent with previous studies showing that root-zone hypoxia constrains mitochondrial respiration and cellular ATP production, thereby limiting metabolically demanding processes such as ion uptake and long-distance transport (Wagner et al., 2018). Likewise, controlled hypoxia experiments using N_2_-based O_2_ displacement have reported altered uptake and partitioning of key minerals, particularly potassium and other ions central to osmotic regulation, processes that are highly sensitive to cellular ATP availability (Jethva et al., 2022).

Mechanistically, these responses are consistent with the expectation that hypoxia-driven ATP limitation reduces plasma-membrane H^+^-ATPase activity, thereby depolarizing the membrane potential and weakening the proton-motive force. Such effects would be expected to constrain voltage-dependent transport pathways, including inward-rectifying K^+^ channels (Moshelion and Moran, 2000), while a reduced ΔpH would diminish the driving force for H^+^-coupled secondary active transport processes (Morsomme and Boutry, 2000). In parallel, the increase in soil pH during N_2_-driven gas displacement may also reflect partial removal of dissolved CO_2_ from the soil solution, superimposed on the metabolic effects of inhibited root respiration. Reduced dissolved CO_2_ would be expected to decrease carbonic acid formation and thereby contribute, at least in part, to the observed pH shift.

Importantly, the reduction in whole-plant transpiration following O_2_ depletion developed progressively over several days (Figure 3A; Supplementary Figure S3A), rather than as an abrupt response. This temporal lag argues against hydraulic failure as a primary consequence of hypoxia and instead supports a gradual decline in root function driven by metabolic impairment. In this framework, reduced respiration under low O_2_ would be expected to weaken the energetic and electrochemical basis for solute uptake, including K^+^ acquisition, thereby reducing the osmotic component of water uptake. Additional energy-dependent constraints, including possible effects on aquaporin function, may further aggravate this decline (Tournaire-Roux et al., 2003; Boursiac et al., 2008). Thus, impaired whole-plant hydraulics are better interpreted here as a downstream consequence of hypoxia-induced metabolic limitation rather than as its immediate cause.

Under waterlogging, whole-plant transpiration in M82 was also reduced, but both the magnitude of this reduction and the associated developmental responses differed from those observed under N_2_-induced hypoxia. Following the onset of waterlogging, transpiration declined rapidly and reached a minimum within approximately 4-5 d, with a mean reduction of about 16% relative to drained controls (Figure 5A; Supplementary Figure S5A). In contrast, N_2_-induced O_2_ displacement caused a somewhat larger decline in transpiration in M82, but this response developed more gradually rather than more rapidly (Figure 3A; Supplementary Figure S3A). This difference suggests that waterlogging and gas-driven hypoxia imposed distinct constraints on root function. N_2_ flushing likely reduced oxygen availability more uniformly throughout the root zone, whereas under waterlogging oxygen availability likely remained more heterogeneous along the soil profile. This interpretation is consistent with the root distribution patterns observed at harvest (Figures 5G and 6G), which suggest a shift in root presence toward the soil surface under waterlogging, where oxygen availability may have been less restricted.

Notably, only a subset of waterlogged M82 plants displayed a pronounced transpiration decline, and these individuals were also characterized by stronger adventitious-root formation (Figures 5A and 5H). Across individuals, adventitious-root emergence was associated with greater transpiration reduction (Figure 5I), indicating that adventitious rooting tracked stress severity rather than representing an early or uniformly protective response. This pattern was modest in M82 but much more pronounced in the introgression lines, in which waterlogging caused substantially larger transpiration reductions, approximately 44% in IL11-4 and 55% in IL8-1 relative to controls (Figure 5A-C; Supplementary Figure S5A-C). Thus, the genotypes differed markedly in their ability to maintain whole-plant water flux under waterlogging. M82 exhibited relatively small transpiration reductions, low adventitious-root scores (Figure 5H), and cumulative transpiration comparable to drained controls (Figure 6A), while maintaining shoot biomass and reproductive output (Figure 6B; Supplementary Figure S6). In contrast, IL11-4 and IL8-1 showed pronounced and sustained transpiration declines, extensive adventitious-root proliferation, reduced shoot growth, lower shoot water-use efficiency, and clear deterioration of the primary root system (Figures 5H,I, 6B-G, and S5B,C). This contrast raises an important question: why did N_2_-induced hypoxia not trigger adventitious-root formation?

Several, not mutually exclusive, explanations may account for the absence of adventitious-root formation in the N_2_-displacement experiment. First, these experiments were conducted only in M82, a genotype that also exhibited relatively limited adventitious-root formation under waterlogging compared with IL11-4 and especially IL8-1 (Figure 5H). Second, the duration of the N_2_-displacement treatment was shorter than that of the waterlogging experiment and may have been insufficient to allow full development of this response. Third, oxygen depletion in the N_2_-displacement system was imposed rapidly, whereas during waterlogging the decline in rhizosphere O_2_ may have developed more gradually.

Adventitious-root induction may therefore depend not only on the severity of oxygen limitation, but also on genotype, exposure duration, and the temporal dynamics of O_2_ decline. Thus, under the conditions used here, N_2_-induced oxygen deprivation alone did not reproduce adventitious-root formation, but these experiments do not exclude a role for slower or longer hypoxic exposure in triggering this developmental response.

Stem cross-sections taken above the adventitious-root zone further suggested a reduction in the visually distinguishable xylem area in waterlogged IL11-4 and IL8-1 plants (Figure 7), consistent with persistent impairment of the primary hydraulic pathway. Quantitatively, the transient partial recovery of transpiration during days 14-22 under waterlogging, coinciding with adventitious-root emergence and followed by renewed decline after drainage (Figure 5A-C; Supplementary Figure S5A-C, during the period marked by the green dashed lines), indicates that adventitious roots contributed only a limited fraction of whole-plant water uptake, on the order of approximately 15%-20% of daily transpiration.

Together, these results indicate that adventitious rooting represents a compensatory response that develops after substantial impairment of the primary root system, sufficient to provide limited hydraulic support but insufficient to restore intact whole-plant water transport. This temporal coupling between root emergence and partial transpiration recovery supports H2 and indicates that adventitious-root formation was not reproduced by the N_2_-displacement treatment used here. This difference may reflect the more gradual decline in rhizosphere O_2_ during waterlogging, together with additional hydraulic or chemical cues associated with water excess, indicating that the developmental response observed under waterlogging was not reproduced by rapid O_2_ displacement alone.

## Conclusions

In tomato, reduced root-zone oxygen availability appears to be the primary driver of waterlogging injury, whereas genotype-specific differences in hydraulic performance, metabolic capacity, and developmental plasticity shape the extent and form of the whole-plant response. By experimentally separating O_2_ deprivation from water excess, we show that root-zone hypoxia constrains whole-plant transpiration primarily through impaired root metabolism and transport capacity, whereas waterlogging imposes additional conditions not reproduced by rapid O_2_ displacement alone. Under waterlogging, adventitious-root development was associated with a transient partial recovery of transpiration, yet quantitative analysis indicates that this recovery reflects only a limited contribution to whole-plant water uptake, insufficient to restore pre-waterlogging transpiration or growth. Thus, adventitious roots function primarily as a compensatory response that provides limited hydraulic support after substantial impairment of the primary root system, rather than as a functional replacement conferring full waterlogging tolerance. Together, these findings provide a quantitative, time-resolved framework for interpreting waterlogging responses and show that adventitious roots represent a limited, genotype-dependent hydraulic bypass rather than a universal acclimation mechanism.

## Materials and Methods

### Plant material and growth conditions

Tomato plants (*Solanum lycopersicum*) from three lines were used: cv. M82 and the two introgression lines IL11-4 and IL8-1 (Eshed and Zamir, 1995). These lines were previously characterized as differing in whole-plant transpiration under drought, with IL11-4 showing higher transpiration, biomass accumulation, and water-use efficiency than IL8-1, whereas M82 exhibited intermediate responses (Gosa et al., 2022). Seeds were germinated in commercial germination trays and grown for 4 weeks in a commercial substrate (Bental 11 Premium Growing Mix; Tuff-Substrates, Israel). Seedlings were then transplanted into pots, soil columns, or waterlogging units (see below) filled with fine quartz sand (Negev Industrial Minerals Ltd., Israel) and placed on the PlantArray functional-phenotyping platform. To minimize soil-surface evaporation, an ethylene-vinyl acetate (EVA) disc was placed over the sand surface. In all experiments, plants were allowed to establish under daily irrigation adjusted to measured transpiration using the functional-phenotyping platform automatic feedback system, thereby maintaining soil water content near field capacity until roots reached the full column depth. During this establishment phase, the system was operated to verify performance and define baseline physiological parameters.

All experiments were conducted in the iCORE functional-phenotyping greenhouse at the Faculty of Agriculture, Food and Environment, Rehovot, Israel, during three experimental periods: spring 2019, autumn 2019, and autumn 2023. The polycarbonate greenhouse was equipped with cooling pads to regulate temperature while allowing natural environmental fluctuations and preventing temperatures from exceeding 32 °C (Figures 1 and 2A). Air temperature, relative humidity (RH), vapor pressure deficit (VPD), and photosynthetic photon flux density (PPFD) were continuously monitored using a central greenhouse weather station (Supplementary Figure S1).

Continuous whole-plant transpiration was measured using the high-throughput gravimetric functional-phenotyping system PlantArray 3.0 (Plant-DiTech, Israel). Plant weight was recorded at 3-min intervals using load cells, and transpiration was calculated from changes in pot weight over time. Data were processed using SPAC Analytics software (Plant-DiTech).

Detailed descriptions of system operation and validation are provided in Halperin et al. (2017), Yaaran et al. (2019), and Dalal et al. (2020).

### Oxygen deficiency induced by oxygen displacement (non-flood hypoxia)

To isolate the effects of root-zone oxygen deficiency independently of soil water excess, we used a controlled soil-column system in which oxygen was displaced from soil pore space by N_2_ gas, thereby imposing hypoxic conditions while maintaining soil structure and water-holding capacity under regular irrigation at near-field-capacity conditions (Figure 1). In experiment 1 (Exp. 1), the N_2_ treatment was compared with natural oxygen supply from the soil surface. In experiment 2 (Exp. 2), it was compared with displacement of soil air by atmospheric air applied at a flow rate similar to that of N_2_ in the former treatment. Each experiment included 10 independent soil columns (n = 10) constructed from rigid PVC cylinders (internal diameter, 10.6 cm; height, 43 cm). Prior to treatment initiation, plants were acclimated to the system and baseline physiological measurements were established.

Hypoxic conditions were generated by continuously supplying high-purity N_2_ gas (99.999%) through flexible silicone tubing inserted into each column at depths of 22 and 29 cm below the soil surface. In Exp. 2, aerated control columns received atmospheric air through an identical tubing configuration. Gas was dispersed within the soil using an air stone positioned at the end of the tube. To prevent soil desiccation during continuous gas flow, the gas was humidified by bubbling through water in a sealed 1.5-L bottle before entering the columns (Supplementary Figure S2).

Control conditions differed between experiments. In Exp. 1, control plants were maintained under ambient soil aeration without gas injection. In Exp. 2, to control for possible effects of continuous gas flow itself, including CO_2_ stripping, atmospheric air was supplied through the same tubing system using a small aquarium pump at low flow. This control air was also humidified before entering the soil (Supplementary Figure S2). In Exp. 1, N2 flow was applied continuously for 8 days, followed by a 5 days post-treatment period without gas injection. In Exp. 2, N2 or air flow was applied continuously for 7 days.

To enable repeated monitoring along the soil profile, six access ports were installed at fixed depths: three at 12.5 cm and three at 27.5 cm below the soil surface (diameter, 2 cm). The ports were sealed with cable adapters, allowing insertion of sensors and electrodes without external air infiltration. At each depth, a redox (oxidation-reduction potential, ORP) electrode, a pH electrode, and a soil-solution sampling port were installed. An opposing port (diameter, 3 cm) housed a horizontally mounted oxygen sensor in a sealed plastic holder.

### Oxygen, redox, pH, and soil water content measurements

Two KE-50 oxygen sensors (Figaro Engineering Inc., Japan) were installed in each soil column at depths of 12.5 and 27.5 cm. Each sensor was housed in a perforated plastic tube (20 ± 2 holes, approximately 1 mm in diameter), forming an equilibration chamber that allowed direct exchange with soil pore gas. The housing was kept in direct contact with the surrounding sand matrix to ensure equilibration with the soil atmosphere rather than with a sealed headspace. Sensor insertion points were sealed with silicone adhesive sealant to prevent external air intrusion.

Redox potential was measured using wide-band platinum electrodes with Ag/AgCl reference cells (ELH-031, Van London Phoenix Co., USA), installed at the same depths as the oxygen sensors and sealed with PG16 adapters. Soil pH was measured using Ag/AgCl^-^ based pH electrodes (ELH-067, Van London Phoenix Co., USA), installed and sealed at the same depths as the redox electrodes.

In each column, 5TE volumetric water-content sensors (Meter Group, USA) were inserted vertically to continuously measure volumetric water content (VWC), temperature, and electrical conductivity (EC) at the two sensor depths. All sensors were connected to the PlantArray controller or to separate dataloggers for continuous data logging.

### Soil solution characterization

Soil solution samples were collected using syringes connected to Rhizon samplers (Rhizosphere Research Products, Wageningen, The Netherlands) installed at the two sensor depths. Soil solution was sampled 2 d before treatment initiation and several hours after treatment termination. Macro- and micronutrient concentrations were quantified by ICP-AES (ARCOS, SPECTRO GmbH, Kleve, Germany).

### Harvesting and tissue analysis

At the end of the experiment, drainage water was collected from the base of each pot and analyzed for elemental composition by ICP-AES. The youngest fully expanded leaf was harvested, fresh weight was recorded, and the tissue was dried at 60 °C for 4 d and ground to a fine powder. For elemental analysis, 100 mg of dry tissue was digested in concentrated nitric acid, heated, and then treated with perchloric acid. The digests were diluted and analyzed by ICP-AES.

Roots were thoroughly rinsed to remove sand, immediately frozen in liquid nitrogen, and stored until analysis. Root elemental composition was determined using low-vacuum scanning electron microscopy coupled with energy-dispersive X-ray spectroscopy (SEM-EDS; JEOL IT100).

### Waterlogging experiment

In this experiment, only the root zone was flooded; therefore, the term waterlogging is used rather than flooding. No direct manipulation of oxygen availability was applied. Changes in rhizosphere oxygen availability arose solely from waterlogging-induced limitations to gas diffusion. Seeds were germinated and grown as described above. One 3-week-old plant was transplanted into each 5-L waterlogging pot (Plant-DiTech “Flood Control Kit” prototype) and placed on the phenotyping platform (Figure 2A). The experiment included 68 pots arranged in a completely randomized design, with 10-12 biological replicates per genotype and treatment (Figure 2B). Following acclimation under standard irrigation-drainage cycles, waterlogging was imposed on half of the plants by closing the drainage outlet, resulting in flooding of the root zone only (Figure 2C). Waterlogging was maintained for 2 weeks, after which plants were returned to normal irrigation-drainage conditions for a 1-week recovery period prior to harvest (Figure 2D). To estimate the contribution of adventitious roots to whole-plant water uptake, we compared the mean daily transpiration loss relative to genotype-matched drained controls during the partial-compensation period under prolonged waterlogging (days 14-22) with the corresponding loss measured immediately after drainage was restored (day 23), when adventitious roots lost access to the surface water layer. The difference between the values of days 23-22 was taken as the effective contribution of adventitious roots to water uptake. The relative contribution was expressed as a percentage of the initial transpiration loss measured on days 13-14. For the quantitative analysis of transpiration loss in Figure 5, included waterlogged plants comprised all IL8-1 plants exhibiting reduced transpiration, all but one IL11-4 plant showing a reduction, and only three M82 plants displaying a clear decrease during the initial decline window (days 13–14).

### Measurement of agro-morphological parameters

Adventitious roots emerging on the sand surface were scored on a 0-5 scale (0, none visible; 5, surface fully covered). Scoring was performed by a single observer using predefined criteria and applied consistently across all treatments. At harvest, shoot and root length, fresh and dry biomass, and the numbers of flowers and fruits were recorded.

### Stem section preparation and analysis

The aerial stem portion was excised immediately above the adventitious-root zone, when present, and a fresh 10-cm segment was fixed in 70% ethanol. Transverse sections (approximately 100 µm thick) were prepared using a rotary microtome (Leica RM2255; TC-65 disposable blades), stained with 0.1% (w/v) toluidine blue, and imaged under bright-field illumination using an Olympus BX53 microscope equipped with a DP73 camera.

### Statistical analysis

Data were processed using SPAC Analytics (Plant-DiTech, Israel) and JMP Pro 18.0 (SAS Institute). The statistical tests applied to each dataset are specified in the corresponding figure legends.

## Acknowledgements

H.P. and M.M. gratefully acknowledge the funding and support provided by the Azrieli Foundation to H.P. through their International Postdoctoral Fellowship Programme.

## Funding

This project was funded by the Planning and Budgeting Committee of the Council for Higher Education in Israel (VATAT) as part of the program “Israeli Center for Digital Agriculture.” Additional support was provided by an individual postdoctoral fellowship from the Azrieli Foundation awarded to H.P., and by the Israel Ministry of Agriculture and Rural Development (Eugene Kandel Knowledge Centers) through the project “Root of the Matter – The root zone knowledge center for leveraging modern agriculture” (Grant No. 391-15). The funding sources had no role in study design, data collection, data analysis, interpretation of the results, manuscript preparation, or the decision to submit the work for publication.

## Author Contributions

H.P. led the second part of the study, including the waterlogging experiments, adventitious-root analyses, and the flood-system setup, and contributed to data analysis and writing. D.M. led the first part of the study, including the non-flood hypoxia experiments and mineral-uptake analyses, and contributed to experiment setup, data analysis, and writing. D.B. contributed to experimental design, experiment setup, and data analysis. I.N. contributed to conceptual development and scientific discussions. M.M. and M.S. conceived the study, designed the experiments, secured funding, supervised the research, and contributed to data interpretation and manuscript writing. All authors reviewed and approved the final manuscript.

## Conflict of interest

The authors declare no competing financial interests or personal relationships that could have influenced the work reported in this paper.

## Data Availability

The datasets generated and/or analyzed during the current study are available from the corresponding authors upon reasonable request.

## Notes

### Competing Interest Statement

The authors have declared no competing interest.

## References

Aslam A, Mahmood A, Ur-Rehman H, Li C, Liang X, Shao J, Negm S, Moustafa M, Aamer M, Hassan MU (2023) Plant Adaptation to Flooding Stress under Changing Climate Conditions: Ongoing Breakthroughs and Future Challenges. Plants 12: 3824 10.3390/plants12223824

Bansal R, Srivastava JP (2015) Effect of waterlogging on photosynthetic and biochemical parameters in pigeonpea. Russ J Plant Physiol 62: 322–327 10.1134/S1021443715030036

Boursiac Y, Boudet J, Postaire O, Luu D, Tournaire-Roux C, Maurel C (2008) Stimulus-induced downregulation of root water transport involves reactive oxygen species-activated cell signalling and plasma membrane intrinsic protein internalization. The Plant Journal 56: 207–218 10.1111/j.1365-313X.2008.03594.x

Colmer TD, Voesenek L (2009) Flooding tolerance: suites of plant traits in variable environments. Functional Plant Biology 36: 665–681 10.1071/FP09144

Dalal A, Shenhar I, Bourstein R, Mayo A, Grunwald Y, Averbuch N, Attia Z, Wallach R, Moshelion M (2020) A telemetric, gravimetric platform for real-time physiological phenotyping of plant–environment interactions. JoVE (Journal of Visualized Experiments) e61280 10.3791/61280

Eshed Y, Zamir D (1995) An introgression line population of Lycopersicon pennellii in the cultivated tomato enables the identification and fine mapping of yield-associated QTL. Genetics 141: 1147–1162 10.1093/genetics/141.3.1147

FAO (2015) The impact of natural hazards and disasters on agriculture. FAO

Gosa SC, Gebeyo BA, Patil R, Mencia R, Moshelion M (2022) Diurnal stomatal apertures profile and density ratios affect whole-canopy conductance, drought response, water-use efficiency and yield. bioRxiv 2022–01 10.1101/2022.01.06.475121

Halperin O, Gebremedhin A, Wallach R, Moshelion M (2017) High-throughput physiological phenotyping and screening system for the characterization of plant– environment interactions. The Plant Journal 89: 839–850 10.1111/tpj.13425

Horchani F, Khayati H, Raymond P, Brouquisse R, Aschi-Smiti S (2009) Contrasted Effects of Prolonged Root Hypoxia on Tomato Root and Fruit (*Solanum lycopersicum*) Metabolism. J Agronomy Crop Science 195: 313–318 10.1111/j.1439-037X.2009.00363.x

Jackson WT (1956) The relative importance of factors causing injury to shoots of flooded tomato plants. American Journal of Botany 637–639 10.2307/2438827

Jethva J, Schmidt RR, Sauter M, Selinski J (2022) Try or die: Dynamics of plant respiration and how to survive low oxygen conditions. Plants 11: 205 10.3390/plants11020205

Jia W, Ma M, Chen J, Wu S (2021) Plant morphological, physiological and anatomical adaption to flooding stress and the underlying molecular mechanisms. International Journal of Molecular Sciences 22: 1088 10.3390/ijms22031088

Kęska K, Szcześniak MW, Makałowska I, Czernicka M (2021) Long-term waterlogging as factor contributing to hypoxia stress tolerance enhancement in cucumber: comparative transcriptome analysis of waterlogging sensitive and tolerant accessions. Genes 12: 189 10.3390/genes12020189

Kuo CG, Chen BW (1980) Physiological Responses of Tomato Cultivars to Flooding. Journal of the American Society for Horticultural Science 105: 751–755 10.21273/JASHS.105.5.751

Leyshon AJ, Sheard RW (1974) Influence of short-term flooding on the growth and plant nutrient composition of barley. Can J Soil Sci 54: 463–473 10.4141/cjss74-060

McNamara ST, Mitchell CA (1990) Adaptive stem and adventitious root responses of two tomato genotypes to flooding. HortScience 25: 100–103 10.21273/HORTSCI.25.1.100

Morsomme P, Boutry M (2000) The plant plasma membrane H+-ATPase: structure, function and regulation. Biochimica et Biophysica Acta (BBA)-Biomembranes 1465: 1–16 10.1016/S0005-2736(00)00128-0

Moshelion M, Moran N (2000) Potassium-efflux channels in extensor and flexor cells of the motor organ of Samanea saman are not identical. Effects of cytosolic calcium. Plant Physiology 124: 911–919 10.1104/pp.124.2.911

Pan J, Sharif R, Xu X, Chen X (2021) Mechanisms of waterlogging tolerance in plants: Research progress and prospects. Frontiers in Plant Science 11: 627331 10.3389/fpls.2020.627331

Pezeshki SR, DeLaune RD (2012) Soil oxidation-reduction in wetlands and its impact on plant functioning. Biology 1: 196–221 10.3390/biology1020196

Rentschler J, Salhab M, Jafino BA (2022) Flood exposure and poverty in 188 countries. Nature communications 13: 3527 10.1038/s41467-022-30727-4

Sauter M (2013) Root responses to flooding. Current Opinion in Plant Biology 16: 282–286 10.1016/j.pbi.2013.03.013

Sauter M, Steffens B (2014) Biogenesis of Adventitious Roots and Their Involvement in the Adaptation to Oxygen Limitations. In JT Van Dongen, F Licausi, eds, Low-Oxygen Stress in Plants. Springer Vienna, Vienna, pp 299–312 10.1007/978-3-7091-1254-0_15

Singh AK, Vijai P, Srivastava JP (2018) Plants under waterlogged conditions: an overview. Engineering practices for Management of Soil Salinity 335–376 10.1201/9781351171083-25

Tournaire-Roux C, Sutka M, Javot H, Gout E, Gerbeau P, Luu D-T, Bligny R, Maurel C (2003) Cytosolic pH regulates root water transport during anoxic stress through gating of aquaporins. Nature 425: 393–397 10.1038/nature01853

Vidoz ML, Loreti E, Mensuali A, Alpi A, Perata P (2010) Hormonal interplay during adventitious root formation in flooded tomato plants. The Plant Journal 63: 551–562 10.1111/j.1365-313X.2010.04262.x

Voesenek LACJ, Bailey-Serres J (2015) Flood adaptive traits and processes: an overview. New Phytologist 206: 57–73 10.1111/nph.13209

Wagner S, Van Aken O, Elsässer M, Schwarzländer M (2018) Mitochondrial energy signaling and its role in the low-oxygen stress response of plants. Plant Physiology 176: 1156–1170 10.1104/pp.17.01387

Yaaran A, Negin B, Moshelion M (2019) Role of guard-cell ABA in determining steady-state stomatal aperture and prompt vapor-pressure-deficit response. Plant science 281: 31–40 10.1016/j.plantsci.2018.12.027

Yalin D, Schwartz A, Tarchitzky J, Shenker M (2021) Soil oxygen and water dynamics underlying hypoxic conditions in the root-zone of avocado irrigated with treated wastewater in clay soil. Soil and Tillage Research 212: 105039 10.1016/j.agwat.2021.107050

Zhang M, Zhai G, He T, Wu C (2023) A growing global threat: Long-term trends show cropland exposure to flooding on the rise. Science of The Total Environment 899: 165675 10.1016/j.scitotenv.2023.165675

