## Supplementary Information for "Adventitious roots facilitate surface water uptake but only partially sustain transpiration under waterlogging in tomato (*Solanum lycopersicum*)"

**Supplementary Table S1**: **The concentration of nutrients in the irrigation solution in which the plants were watered throughout the experiment.**

| **Element** | **Concentration [ppm]** |
| --- | --- |
| N-NO_3_ | 69.9 |
| P | 30 |
| K | 180 |
| S | 82.2 |
| Mg | 61.4 |
| *Fe | 2.71 |
| Mn | 1.79 |
| Zn | 0.300 |
| Cu | 0.126 |
| B | 0.034 |
| Mo | 0.087 |

* Fe - Given in the irrigation solution in complex with EDTA .

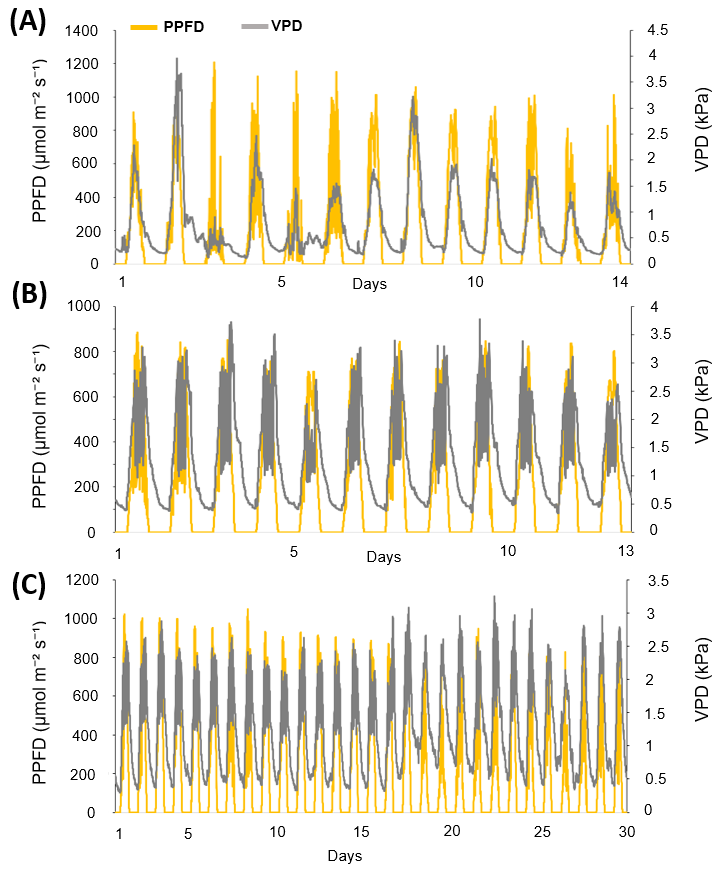

**Supplementary Figure S1: Photosynthetic photon flux density (PPFD) and ambient vapor pressure deficit (VPD) during the experiments.** (A) Spring 2019; (B) Autumn 2019; (C) Autumn 2023. Measurements were recorded continuously by the in-greenhouse weather station; PPFD and VPD were monitored at the greenhouse center. Left axis: PPFD (µmol m⁻² s⁻¹). Right axis: VPD (kPa).

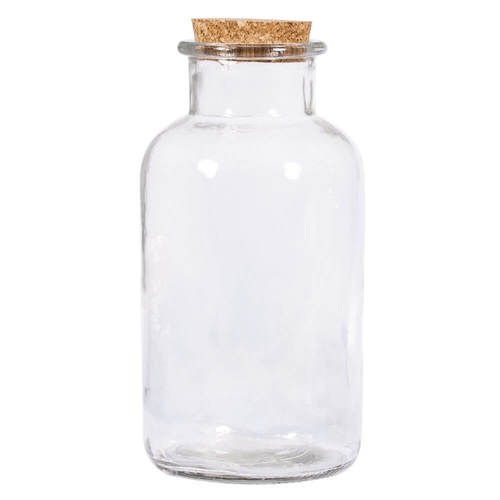

air or nitrogen

saturated air or nitrogen

**Supplementary Figure S2. Gas-humidification bottle for hypoxia treatments.** A closed 1.5 L bottle humidifies the gas: the long inlet (immersed) brings N₂ or air into the water, bubbles rise and humidify, and the short outlet (not immersed) carries saturated N₂ or air to the root.

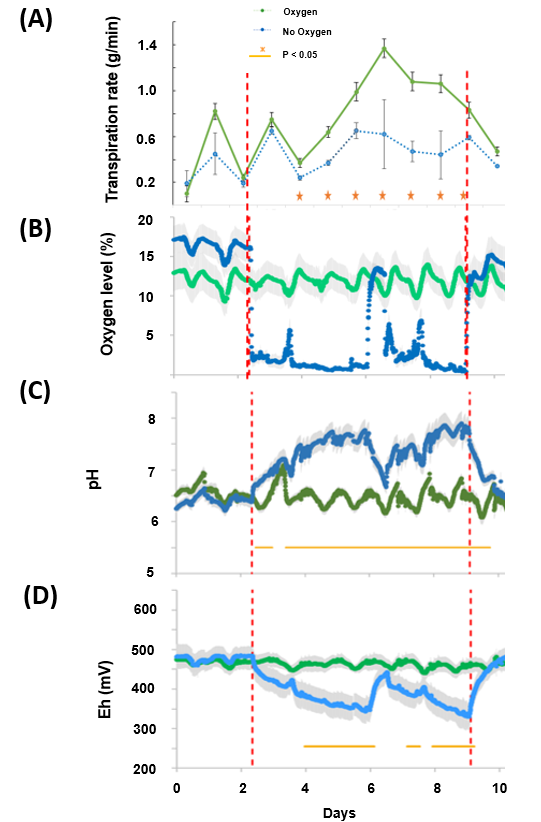

**Supplementary Figure S3. Time series of key measurements in experiment 1.** (A) Whole-plant transpiration rate at midday (mean ± SE; n = 5). (B) Root-zone oxygen level (mean ± SE; n = 10). (C) pH (mean ± SE; n = 10). (D) Redox potential (Eh; mean ± SE; n = 10). Red dashed lines mark the start and end of nitrogen injection (oxygen-deficiency treatment). Orange symbols/lines indicate time points or intervals with significant differences between treatments (p < 0.05). A brief interruption in nitrogen injection occurred on day 6, causing transient deviations. After eight days of treatment, plants remained for five additional days without gas injection; this post-treatment period is included in the time series.

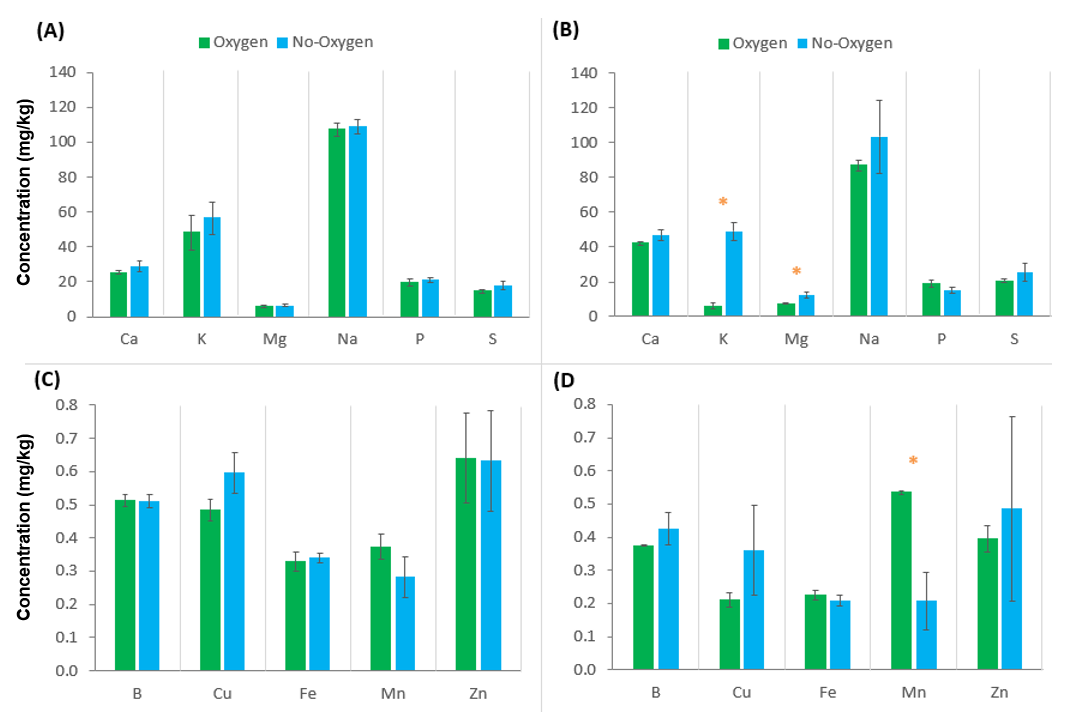

**Supplementary Figure S4. Soil-solution elemental concentrations before and after hypoxia (experiment 1).** (A,B) Macroelements; (C,D) microelements. Panels A and C: two days before treatment. Panels B and D: several hours after treatment cessation. Bars are mean ± SE. Sample sizes: A,C—Oxygen n = 5, No-oxygen n = 4; B,D—n = 5. Units: mg L⁻¹. Green = ventilated control; blue = no-oxygen. Orange asterisks indicate significant differences between treatments (t-test, p < 0.05).

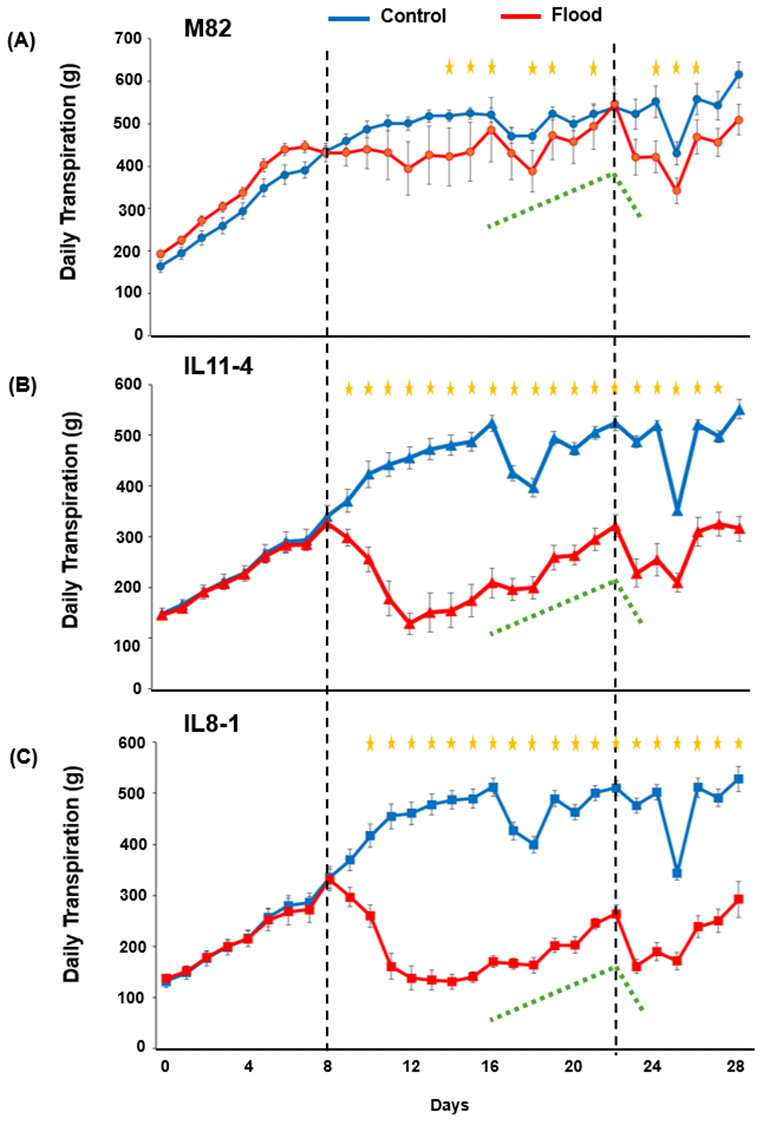

**Supplementary Figure S5. Daily transpiration dynamics of tomato genotypes under waterlogging.** Daily whole-plant transpiration of (A) M82, (B) IL11-4, and (C) IL8-1 grown under control (blue) or waterlogging (red) conditions. Data are presented as means ± SE. Vertical dashed lines indicate the onset and termination of the waterlogging treatment. Orange asterisks denote days with statistically significant differences between treatments (p < 0.05). Dotted green lines highlight the period of partial transpiration recovery observed during prolonged waterlogging, followed by a renewed decline after drainage and return to irrigation–drainage cycles.

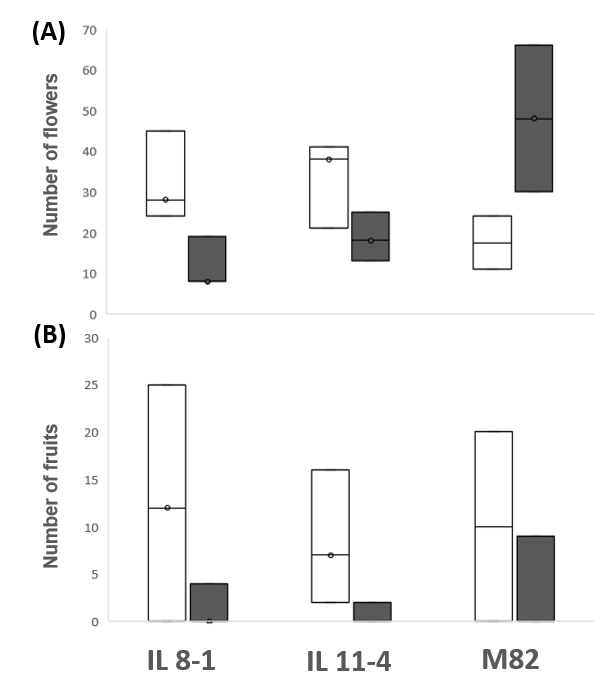

**Supplementary Figure S6. Effects of waterlogging on reproductive development in tomato (Experiment 3).** (A) Number of flowers and (B) number of fruits per plant across genotypes (IL8-1, IL11-4, M82) under control and waterlogging conditions. Boxplots show median, interquartile range, and outliers (n = 5 per treatment). White boxes represent control plants; filled boxes represent waterlogged plants.
